# Characterization of predictive power of extracellular signal recordings in a cerebral ischemia animal model

**DOI:** 10.1101/2023.05.19.541417

**Authors:** S. Matinyan, L. Hovhannisyan, A. Hakobyan, L. Hayrapetyan, A. Khamperyan, K. Yenkoyan

## Abstract

**Introduction:** Global cerebral ischemia leads to substantial and irreversible damage of brain tissue. As it progresses in a less severe course, certain strategies should be implemented to screen this condition on its early onset. The research aimed to characterize the local field potential (LFP) alterations recorded during the subacute phase of the cerebral ischemia animal model and provide their predictive power.

**Methods:** The extracellular signal recordings from the parietal cortex of animals were registered with a neural probe. The signal was amplified, filtered, digitized, and acquired with Intan amplifier and USB interface boards. The recordings were obtained both in normal conditions and after implementation of unilateral common carotid artery occlusion. The data analysis and classification were performed using NI Diadem software and custom-written code in IPython environment. The respective morphological changes were screened in cerebral cortex and hippocampus. The whole-brain slicing and TTC staining were used for infarct size evaluation.

**Results:** In Fourier spectrograms of intact brain recordings, a peak at 14.4-15 Hz frequencies was detected, whereas this phenomenon was absent in cerebral ischemia model recordings. In channels’ cross-correlograms for intact and ischemic brain recordings, there was a clear difference detected in the maximum peak power. With autocorrelation analysis, the long lag rhythmicity was detected in normal brain recordings, while no rhythmicity was observed in ischemic brains. The morphological and behavioral analyses did not result in any significant changes and neural loss. The TTC staining failed to show any damaged area ipsilateral to the occluded common carotid artery.

**Conclusion:** We have analyzed and described the major characteristics of the electrical activity that vary between neural populations of the parietal cortex of normal and ischemic brains. This data proves that LFP recordings can be used for further investigation of changes occurring in the subacute phase after unilateral common carotid artery occlusion.

## 1. Introduction

Ischemic injury as a result of the global circulatory arrest in the brain is one of the major causes of death and disability in the adult population (Du et al., 2017). Although many studies are available concerning further alterations following acute focal cerebral ischemia, no consistent data is available concerning the damage induced by chronic cerebral ischemia (CCH). CCH is characterized by a gradual memory loss, progressive cognitive decline, and alterations in behavior and social cognition (Series & Esiri, 2012). The clinical sign of CCH, namely vascular cognitive impairment and dementia (VCID) is the second most prevalent cause of cognitive impairment after Alzheimer’s disease (Yoshizaki et al., 2008). It appears in about 20% of all dementia patients and this is predicted to triple by 2050. Therefore, it has excessive clinical and social impact and requires significant efforts from scientists and clinicians to implement suitable preventive measures. As this phenomenon flows in a less severe course, certain “markers” should be found to detect this condition from its very onset. The determination of the unique features caused by CCH may also be helpful in diagnosis as well as in testing treatment modalities. The current pharmacological and non-pharmacological approaches are not efficient enough and further studies are required to improve the outcome of applied treatments. The diagnosis of this pathology by itself also represents a huge challenge among clinicians, although the explicit history taking from both the patient and the close relative or friend, physical examination, mental evaluation, multiple questionnaires, biochemical tests, and neuroimaging together can relieve the existing problem (Arvanitakis et al., 2019). The brain circuits work in a precisely defined spatiotemporal pattern and are highly sensitive to minimal fluctuations that come into play during cerebral ischemia due to glucose and oxygen deficiency. In this context electrophysiological recordings can become a sensitive tool for recording of neuronal activity owing to their high temporal resolution, usability in both acute and chronic experiments and possibility to increase the informativeness by multisite recordings of brain regions (Haumesser et al., 2017). From this perspective, the damage caused by ischemic events may have its fingerprints in the electrophysiological activity such as local field potentials recordings (LFPs), that are believed to show the complex, synchronized activity of neural populations. This method has been applied in studies searching for neural alterations in epilepsy, local ischemic injuries, Parkinson’s disease, and hyperkinetic diseases such as dystonia and Huntington’s disease (Wu et al., 2013; S. Wang, Zhang, Sheng, Lu, & Miao, 2015; M. Wang et al., 2018; Zhu et al., 2018). Whether this method can be useful for the early prediction of ischemic events remains unclear.

In this paper, we aim to show, that the LFP recordings can be potentially used as a data supplier for the early detection of the neural activity shifts during cerebral ischemia.

## 2. Materials and methods

### 2.1. Unilateral common carotid artery occlusion model

To describe the recording results from the ipsilateral parietal cortex right unilateral common carotid artery occlusion (UCCAO) procedure (n = 9) was performed (Venkat et al., 2015). The rats were anesthetized with pentobarbital (50 mg/kg). Ketoprofen (5 mg/kg) was injected to prevent post-surgical pain. By midline cervical incision the right common carotid artery was isolated, separated from the vagus nerve double-ligated with 5.0 sutures (Banoujaafar et al., 2014). The artery was cut between the sutures to ensure the completeness of the procedure. The sham-operated animals (n = 9) received the same procedure without common carotid artery ligation.

### 2.1. Surgical procedure for electrode implantation

Just after the procedure, the animals were put on the stereotaxic equipment. The bregma-lambda distance has been measured and the stereotactic coordinates have been calculated according to the atlas (G.Paxinos & Ch.Watson, 2007). Lidocaine spray has been applied locally and dexamethasone (0.2mg/kg) has been injected i/m to prevent brain swelling. A window of 4mm*6mm has been drilled and the electrode has been slowly inserted in AP (0-4.9mm), ML (1.6mm), DV (1.5mm) coordinates, i.e. the electrode comb having length of 5mm has been inserted parallel to sagittal suture. The reference wire was placed on the scalp. The probe has been fixed by dental acrylic and the animals were left to recover for 7 days before the recordings. The animals were kept under standard laboratory conditions with 12h light regimen (7:00-19:00) with full access to water and food. All experiments were conducted according to European Communities Council Directive (86/609/EEC) and were approved by the institutional ethical committee.

### 2.3. Electrophysiological recordings

The details of neural probe fabrication are described in Muradyan et al. (S.Muradyan, A.Hakobyan, 2018). The neural probe was fabricated according to the following criteria: to match the size of the chosen rat brain, to maintain the appropriate proportion of the recording site, and to minimize the implantation-induced brain injury. The designed probes had 5 electrode tips that are 100μm wide and in a 5mm long array separations between tips are 600 – 800μm. The impedance of recording sites against reference wire was measured in saline solution and estimated to be 6kΩ – 8kΩ at a 1KHz signal frequency. The recording experiments were started after 7 days of postsurgical recovery period and performed under pentobarbital/xylazine anesthesia in the grounded Faraday cage. The electrical activity was amplified, notch filtered and digitized at a 20 KHz sampling rate with Intan RHD2000 series amplifier board and acquired via Intan Technologies USB interface Board and open-source recording program.

### 2.4. Morphological staining

#### 2.4.1. Hematoxylin-eosin staining

The animals (n = 3 in control and experimental groups) were anesthetized with pentobarbital (10 mg/kg body weight via intraperitoneal injection, 3∼5 min before perfusion) on the 7^th^ day after surgery. Then they were perfused with 0.01 M PBS solution and 4% paraformaldehyde solution. The brain tissue was separated from the rats’ corpse and then placed into 4% paraformaldehyde solution for 18 h. The H & E staining was performed as described by Y.Li et al (Y. Li et al., 2018). Briefly, after keeping the tissues in Harris’ hematoxylin solution for 6 h, they were rinsed without and with 10% acetic acid and 85% ethanol containing tap water. As a final step, the samples were kept in eosin Y ethanol solution for 48 h. After paraffin embedding the brain tissues were cut into 4-μm slices, dewaxed, and mounted with mounting media.

#### 2.4.2. Luxol Fast Blue staining

The brain slices (n = 3 in each group) were stained with Luxol Fast Blue staining according to recommendations(Lockard & Reers, 2009). The formalin-fixed slices were stained at 57° C for 2 h, then at 22° C for 30 min, in a 0.0125% solution of Luxol fast blue in 95% alcohol acidified by 0.1% acetic acid. The sections were washed in distilled water and then treated with a clearing solution.

#### 2.4.3. 2,3,5-triphenyltetrazolium chloride (TTC) staining

The animals (n = 3 in each group) were sacrificed according to the experimental design on the 7^th^ day for stroke size evaluation. The perfused brains were removed quickly and frozen at 0 °C refrigerator for 10 min. Then the brains were dissected to 2 mm slices and incubated for 10 minutes in freshly prepared 0.5% TTC solution (2, 3, 5-Triphenyltetrazolium chloride, Sigma-Aldrich). Afterward, the sections were transferred to 4% paraformaldehyde (L. Li et al., 2018). The slices were photographed and analyzed by ImageJ software (National Institutes of Health, USA) (*ImageJ*, n.d.).

### 2.5. Behavioral testing

#### 2.5.1. Y maze

For the evaluation of spontaneous alternation and locomotor activity Y-shaped maze (n = 9 in each group) has been used as described in (Melkonyan et al., 2021). The animal was placed on the center and was allowed to freely explore the three arms for 5 min. Each trial was recorded by the camera placed 2.0 m above the apparatus. After each trial the maze was cleaned with 70% ethanol solution and allowed to dry. An entry was calculated when the animal crossed the edge of the respective arm with all four limbs. The alternation was considered as a consecutive entry to three different arms. Different sequences were calculated and the percentage of alternation was calculated with the following formula: ((number of alternations)/(total number of arm entries—2) × 100).

#### 2.5.2. Novel object recognition task

Novel object recognition task was conducted in a chamber with plastic walls and back plastic floor. The room was dark during the test period and only the same person was allowed to perform the trials. On the day before the animals were placed to habituate to the empty box for 1 h. The next day two trials were performed. In the first trial, the animals were allowed to explore two identical objects placed in the opposite corners of the box for 10 min. After 1 h break the animals were put back into the box with the replaced novel object for the 5 min period of exploration. The behavior of rats was video recorded. The time spent with novel object and the discrimination ratio (%; time spent in exploring the new object − time spent in exploring the same object)/(time spent in exploring the new object + time spent in exploring the same object) × 100% were calculated (Pan et al., 2017).

### 2.5. Data analysis

The processing of raw data measurement, frequency domain representation and analysis were done in NI DIAdem software environment. Further steps of analysis and construction of classification model were done with custom written code based on Scikit-learn library in IPython environment. To extract features for classification, a short-time Fourier transform with given number of intervals and overlapping segments was applied to raw channel data. The output dataset was composed of 60 features and 20000 samples for each channel. As a feature selection and data reduction, step principal component analysis (PCA) with 30 components was performed on a feature dataset, and the output of PCA was fed to a non-linear support vector machine (SVM) classifier. The model performance was assessed with the accuracy metric. P values <0.05 were considered as statistically significant.

## 3. Results

### 3.1. Morphological findings

The cortical and hippocampal slices in control and experimental animals are available in Figures 1 and 2. The common carotid artery occlusion failed to bring neural loss as seen in the figures. Cell structures and layers were preserved in cortical areas after the procedure in hematoxylin-eosin stained slices (Figure 1). On the Luxol fast blue stained brain areas, we observed increased astrocytosis and periarterial edema, although it was considered as non-significant between the groups. We observed a relatively similar number of cells per 1mm square. The cells in all cortical layers preserved their characteristic shape. The hippocampus was distinguishable and the cellular structures were visible as well (Figure 2). The TTC staining strengthened the previously obtained results. (Figure 1 I, j).

**Figure 1.**
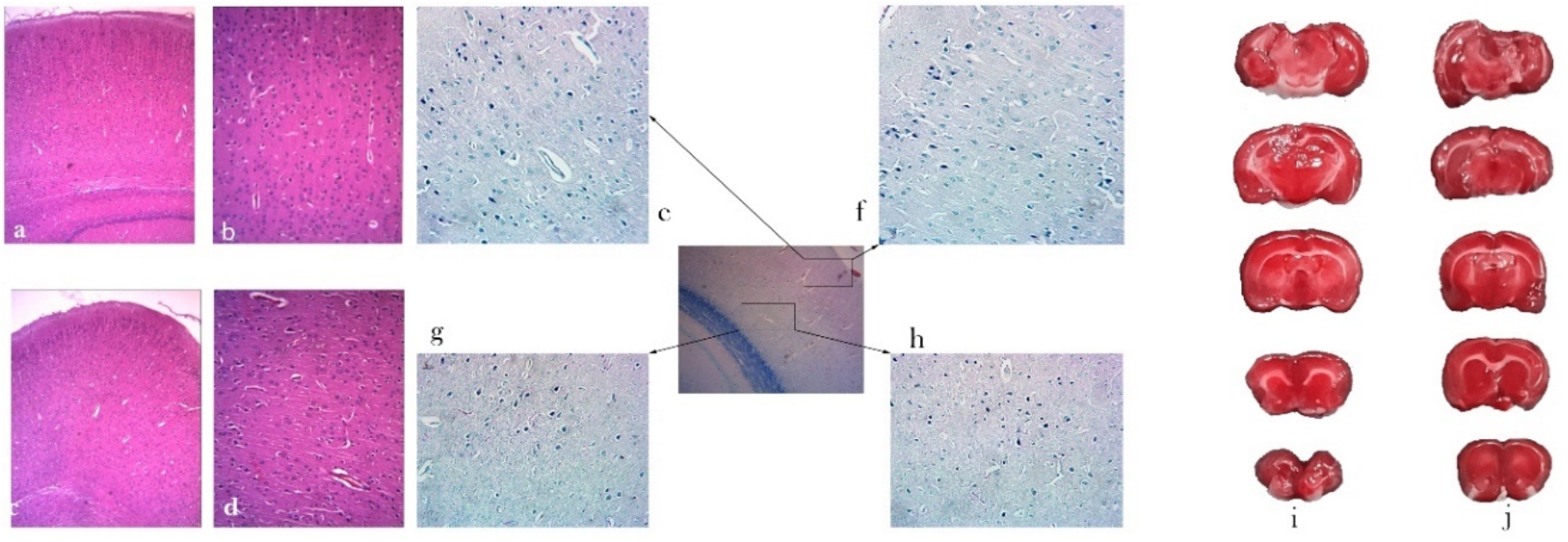
Cortical slices of the animals in sham-operated animals (a, b, e, f) and after common carotid artery occlusion (c, d, g, h). e, f - 1-3 layers of cerebral cortex, g, h-4-6 layers of cerebral cortex. The TTC stained slices in normal (i) conditions and after occlusion of common carotid artery (j). Magnification 5*10 (a, c, 10*10 (b, d), 16*5 (e, f, g, h). a, b, c, d - Hematoxylin-eosin staining; e, f, g, h- Luxol fast blue staining.

**Figure 2.**
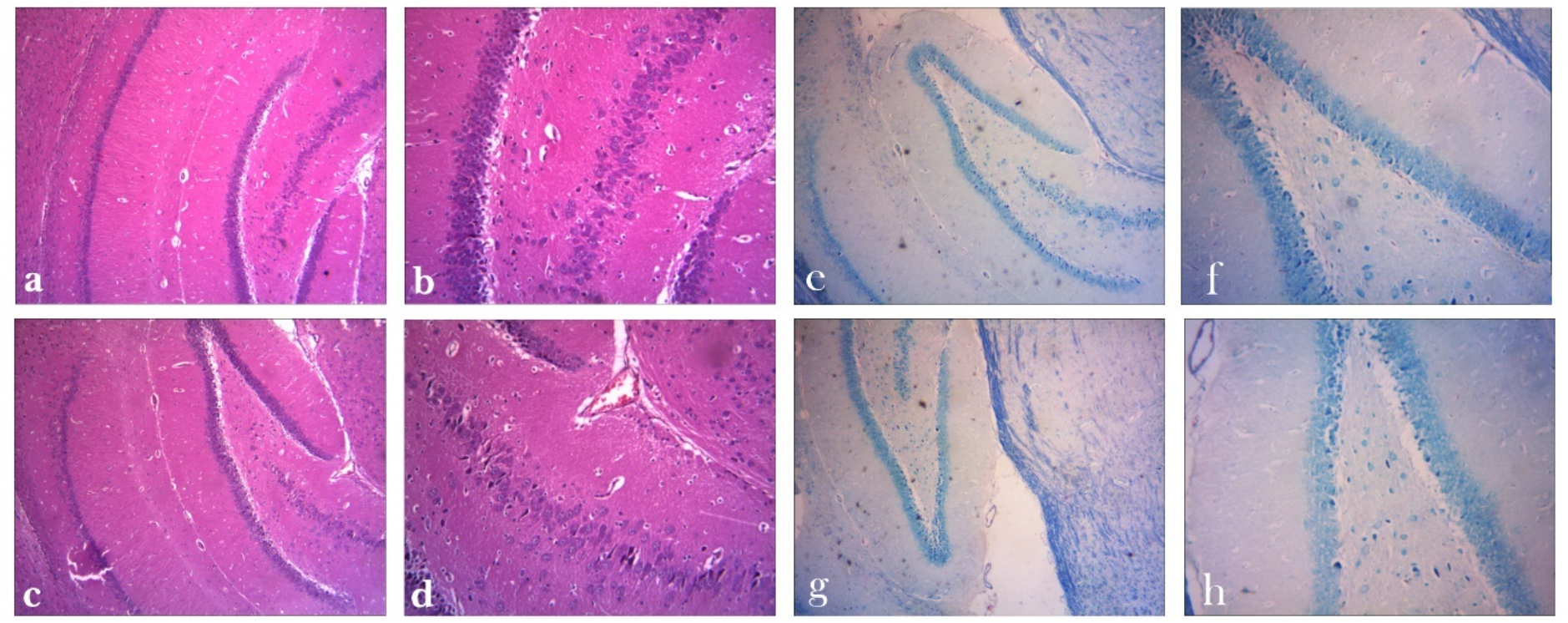
Hippocampal slices of the animals in normal conditions (a, b, e, f) and after common carotid artery occlusion (c, d, g, h). Magnification 5*10 (a, b), 10*10 (b, d), 16*5 (e, f, g, h). a, b, c, d - Hematoxylin-eosin staining; e, f, g, h-Luxol fast blue staining.

### 3.1. Electrophysiological recordings

Although Fourier spectrograms indicate the dominance of slow oscillations in both intact brain and global ischemia model recordings, there were clear differences of power spectra in some frequency ranges in all channels, especially emphasized in 14.4-15 Hz frequencies (Figure 3). In cross spectrograms between channels of each recording the distinction of the maximum peak power can be seen (Fig. 3b). In autocorrelograms summary of 1st windows of LFP recordings the long lag rhythmicity can be observed in intact brain recordings (Fig. 3c) whereas there is no rhythmicity in recording channels of ischemic brain (Fig. 3d).

**Fig 3.**
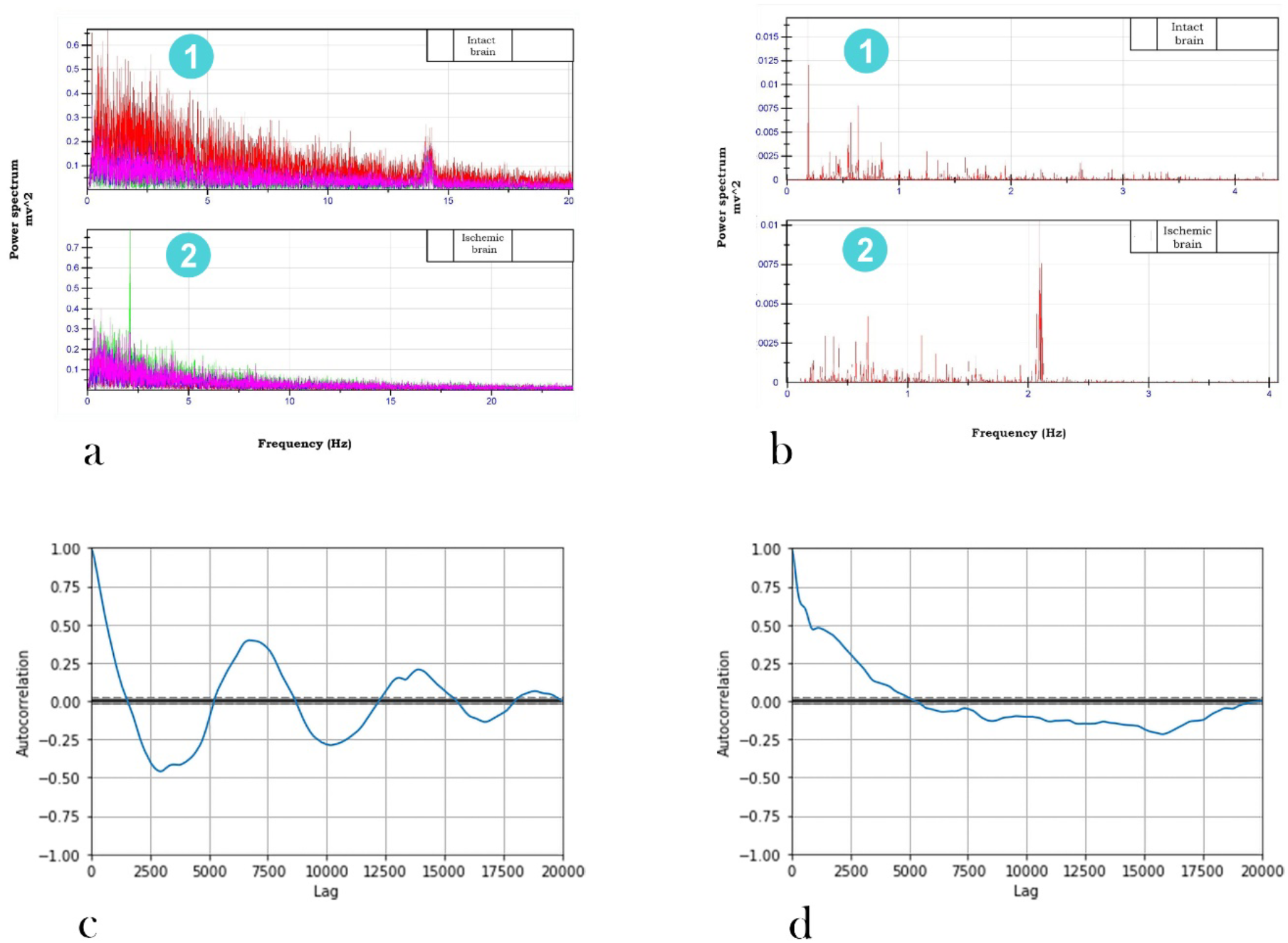
Cumulative spectrograms of LFP for intact (a 1) and ischemic (a 2) brains. Cumulative cross spectrograms of recorded channels for intact (b 1) and ischemic (b 2) brains. Autocorrelogram summary of the intact brain (c) and ischemic brain recording channel (d).

The classification performance of SVC with radial basis function kernel estimated with accuracy metric was 83%. The classification report generated with the classification report function of Scikit-learn library is represented in table 1 (Table 1).

**Table 1.** Classification report.

|  | **Precision** | **Recall** | **F1-score** | **Support** |
| --- | --- | --- | --- | --- |
| Ischemic | 0.77 | 0.92 | 0.84 | 72120 |
| Normal | 0.90 | 0.72 | 0.80 | 71880 |
| Avg/total | 0.83 | 0.82 | 0.82 | 144000 |

### 3.3. Behavioural findings

The spontaneous alternation results (85 ± 11.9 % vs 82± 15.64 %) and the number of entries (9±1 vs 10±4) between sham-operated and experimental animals were not significantly different. We have seen in experimental animals a tendency for more prolonged time spent with the novel object (12.78±4.14 vs 17.83± 6.59) (Figure 4).

**Figure 4.**
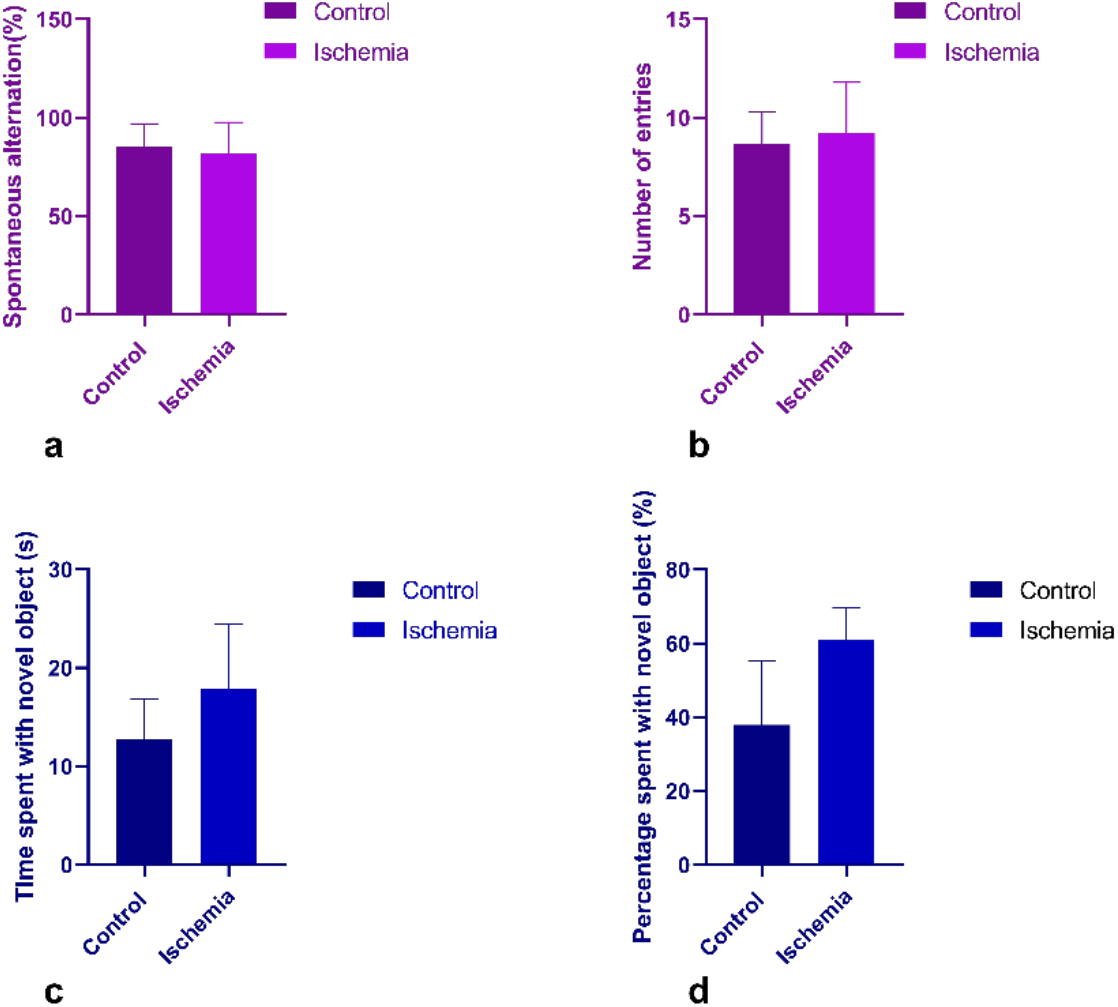
The spontaneous alteration (a) and number of entries (b) between the control and experimental groups in Y maze test. The time spent with novel object (c) and percentage of time spent with novel object (d) between the control and experimental groups in novel object recognition test. Data is presented in mean ± SD form.

## 4. Discussion

The electrophysiological alterations due to ischemic changes has long been a valid routine test for heart diseases with electrocardiogram being among the most popular tests used in clinic (Herring & Paterson, 2006). On the other hand, electroencephalogram (EEG), a non-invasive tool for human brain signal recording, demonstrated certain hallmarks in gamma oscillations, notable as in Alzheimer’s disease (AD) patients, as well as in AD mouse model (Nakazono et al., 2017; Nakazono, Jun, Blurton-Jones, Green, & Igarashi, 2018). Decoding of local field potentials in brain can be promising diagnostic tool from the point of translational medicine and reverse engineering approach in medicine (Telenczuk et al., 2017). They are believed to show the extracellular activity of neural populations and are widely investigated in the context of behavior. What are the main characteristics of extracted data in terms of cerebral ischemia and to what extent they can be predictable? LFP have been used for the assessment of cognitive status in vascular dementia rat model by simultaneous recording from hippocampal CA3 and CA1 regions. Here, electrical oscillations have been subjected to theta-gamma coupling, a method that has been widely used to assess cognitive changes, and have been supported by behavioral and morphological data (Xu et al., 2015). The limitation of this biomarker is the requirement of deep placement of the electrodes and damage to the hippocampus, which would be especially unpleasant as a diagnostic tool for patients already having cognitive issues.

Controversially, LFP, as a representation of neuronal population activity, has been not only attractive information to study particular information flow but also has been subjected to criticism as a concept, being the root of complications for the mathematical modeling and data analysis. Such as, the recorded potentials are not always representing the local events but are affected by global events, or not representing a single population but an overlap of different populations (Herreras, 2016).

In this paper we have hypothesized that LFP can be used for understanding and predicting the neural activity changes caused by unilateral common carotid artery occlusion. We have analyzed the obtained data, which was recorded from the same brain region both under normal conditions and after inducing global ischemia. The data has been analyzed in the following sequence: Short-time Fourier transform (STFT) has been implemented for feature extraction, PCA for feature selection, and non-linear SVM classifier has analyzed the output (STFT->PCA - > SVM).

In Fourier spectrograms, we could clearly observe differences in power spectra, notably in 14.4-15 Hz frequencies. The long lag rhythmicity was absent when recording from the ischemic brain. In addition, the nonlinear SVM classifier model could learn the differences between the two classes. This can be a good basis for future implementation of the tool of preclinical therapeutic interventions assessment in brain recovery processes.

The used morphological approach, although limited, provides solid evidence for the absence of neural loss after artery occlusion, which is in line with previously obtained results (Venkat et al., 2015). Together with LFP changes, these findings suggest disturbances of large-scale neural network activity after the procedure, rather than neural loss. Although having a fingerprint on the neural activity pattern, the animals performed well in behavioral tasks.

In conclusion, this article demonstrates the possibility of using LFP recordings as a diagnostic tool for cerebral ischemia. Future remarks would be the improvement of the current method with more samples and to have stronger evidence for the signatures we have identified. The correctly designed method will play a significant role in the studies dedicated to the investigation of mechanisms and finding therapeutic targets for this disease.

### Limitations of the study

The limitations of the current experiment include the testing on a small subset of animals. Although the brain recordings vary strictly in different animals, which impacts on the average data from different samples, still the environment-related impact requires to have a simultaneous recording of ischemic and non-ischemic samples to exclude their effect, which will be targeted in our future studies.

## 5. Acknowledgment

We would like to thank ISTC Joint Research Grant Program of Innovative Solutions and Technology Center (ISTC) and Enterprise Incubator Foundation (EIF) for supporting our project.

## REFERENCES

Arvanitakis, Z., Shah, R. C., & Bennett, D. A. (2019). Diagnosis and Management of Dementia: Review. JAMA - Journal of the American Medical Association, 322(16), 1589–1599. https://doi.org/10.1001/jama.2019.4782

Banoujaafar, H., Van Hoecke, J., Mossiat, C. M., & Marie, C. (2014). Brain BDNF levels elevation induced by physical training is reduced after unilateral common carotid artery occlusion in rats. Journal of Cerebral Blood Flow and Metabolism : Official Journal of the International Society of Cerebral Blood Flow and Metabolism, 34(10), 1681–1687. https://doi.org/10.1038/jcbfm.2014.133

Du, S. Q., Wang, X. R., Xiao, L. Y., Tu, J. F., Zhu, W., He, T., & Liu, C. Z. (2017). Molecular Mechanisms of Vascular Dementia: What Can Be Learned from Animal Models of Chronic Cerebral Hypoperfusion? Molecular Neurobiology, 54(5), 3670–3682. https://doi.org/10.1007/s12035-016-9915-1

G. Paxinos & Ch. Watson. (2007). The Rat Brain in stereotaxic coordinates (6th editio). Elsevier Inc.

Haumesser, J. K., Kühn, J., Güttler, C., Nguyen, D.-H., Beck, M. H., Kühn, A. A., & van Riesen, C. (2017). Acute & lt;em & gt;In Vivo & lt;/em & gt; Electrophysiological Recordings of Local Field Potentials and Multi-unit Activity from the Hyperdirect Pathway in Anesthetized Rats. Journal of Visualized Experiments, 124, e55940. https://doi.org/10.3791/55940

Herreras, O. (2016). Local field potentials: Myths and misunderstandings. Frontiers in Neural Circuits, 10(DEC), 1–16. https://doi.org/10.3389/fncir.2016.00101

Herring, N., & Paterson, D. J. (2006). ECG diagnosis of acute ischaemia and infarction: past, present and future. QJM, 99(4), 219–230. https://doi.org/10.1093/qjmed/hcl025

ImageJ. (n.d.). Retrieved January 12, 2022, from https://imagej.nih.gov/ij/

Li, L., Yu, Q., & Liang, W. (2018). Use of 2,3,5-triphenyltetrazolium chloride-stained brain tissues for immunofluorescence analyses after focal cerebral ischemia in rats. Pathology, Research and Practice, 214(1), 174–179. https://doi.org/10.1016/J.PRP.2017.11.016

Li, Y., Li, N., Yu, X., Huang, K., Zheng, T., Cheng, X., Zeng, S., & Liu, X. (2018). Hematoxylin and eosin staining of intact tissues via delipidation and ultrasound. Scientific Reports, 8(1), 12259. https://doi.org/10.1038/s41598-018-30755-5

Lockard, I., & Reers, B. L. (2009). Staining Tissue of the Central Nervous System with Luxol Fast Blue and Neutral Red. Http://Dx.Doi.Org/10.3109/10520296209114562, 37(1), 13–16. https://doi.org/10.3109/10520296209114562

Melkonyan, M., Manukyan, A., Hunanyan, L., Grigoryan, A., Harutyunyan, H., Sukiasyan, L., Danielyan, L., & Yenkoyan, K. (2021). Alpha2-Adrenoblockers Regulate Development of Oxidative Stress and Cognitive Behaviour of Rats under Chronic Acoustic Stress Conditions. Pharmaceuticals 2021, Vol. 14, Page 529, 14(6), 529. https://doi.org/10.3390/PH14060529

Nakazono, T., Jun, H., Blurton-Jones, M., Green, K. N., & Igarashi, K. M. (2018). Gamma oscillations in the entorhinal-hippocampal circuit underlying memory and dementia. Neuroscience Research, 129, 40–46. https://doi.org/10.1016/j.neures.2018.02.002

Nakazono, T., Lam, T. N., Patel, A. Y., Kitazawa, M., Saito, T., Saido, T. C., & Igarashi, K. M. (2017). Impaired in vivo gamma oscillations in the medial entorhinal cortex of knock-in Alzheimer model. Frontiers in Systems Neuroscience, 11(June), 1–12. https://doi.org/10.3389/fnsys.2017.00048

Pan, X., Jiang, T., Zhang, L., Zheng, H., Luo, J., & Hu, X. (2017). Physical exercise promotes novel object recognition memory in spontaneously hypertensive rats after ischemic stroke by promoting neural plasticity in the entorhinal cortex. Frontiers in Behavioral Neuroscience, 11, 185. https://doi.org/10.3389/FNBEH.2017.00185/BIBTEX

S. Muradyan, A. Hakobyan, L. H. (2018). Sensors & Transducers Low-cost Electrode Arrays for Recordings of Neural Activity. Sensors & Tranducers, 220(2), 45–50.

Series, H., & Esiri, M. (2012). Vascular dementia: a pragmatic review. Advances in Psychiatric Treatment, 18(5), 372–380. https://doi.org/10.1192/apt.bp.110.008888

Telenczuk, B., Dehghani, N., Le Van Quyen, M., Cash, S. S., Halgren, E., Hatsopoulos, N. G., & Destexhe, A. (2017). Local field potentials primarily reflect inhibitory neuron activity in human and monkey cortex. Scientific Reports, 7(December 2016), 1–10. https://doi.org/10.1038/srep40211

Venkat, P., Chopp, M., & Chen, J. (2015). Models and mechanisms of vascular dementia. Experimental Neurology, 272, 97–108. https://doi.org/10.1016/J.EXPNEUROL.2015.05.006

Wang, M., Song, Y., Zhang, S., Xu, S., Xiao, G., Li, Z., Gao, F., Zhang, Y., Yue, F., Chan, P., & Cai, X. (2018). Abnormal Spontaneous Neuronal Discharge and Local Field Potential both in Cortex and Striatum of a Nonhuman Primate of Parkinson’s Disease using Implantable Microelectrode Arrays. Proceedings of the Annual International Conference of the IEEE Engineering in Medicine and Biology Society, EMBS, 2018-July, 3438–3441. https://doi.org/10.1109/EMBC.2018.8512999

Wang, S., Zhang, J., Sheng, T., Lu, W., & Miao, D. (2015). Hippocampal ischemia causes deficits in local field potential and synaptic plasticity. The Journal of Biomedical Research, 29(March), 370–379. https://doi.org/10.7555/JBR.29.20150010

Wu, J. X., Yang, H., Peng, Y. F., Fang, L. J., Zheng, W., & Song, Z. (2013). The role of local field potential coupling in epileptic synchronization. Neural Regeneration Research, 8(8), 745–753. https://doi.org/10.3969/j.issn.1673-5374.2013.08.009

Xu, X., Liu, C., Li, Z., & Zhang, T. (2015). Effects of Hydrogen Sulfide on Modulation of Theta–Gamma Coupling in Hippocampus in Vascular Dementia Rats. Brain Topography, 28(6), 879–894. https://doi.org/10.1007/s10548-015-0430-x

Yoshizaki, K., Adachi, K., Kataoka, S., Watanabe, A., Tabira, T., Takahashi, K., & Wakita, H. (2008). Chronic cerebral hypoperfusion induced by right unilateral common carotid artery occlusion causes delayed white matter lesions and cognitive impairment in adult mice. Experimental Neurology, 210(2), 585–591. https://doi.org/10.1016/j.expneurol.2007.12.005

Zhu, G., Geng, X., Tan, Z., Chen, Y., Zhang, R., Wang, X., Aziz, T., Wang, S., & Zhang, J. (2018). Characteristics of globus pallidus internus local field potentials in hyperkinetic disease. Frontiers in Neurology, 9(NOV), 1–9. https://doi.org/10.3389/fneur.2018.00934

